# Predator-secreted sulfolipids induce fear-like defense responses in *C. elegans*

**DOI:** 10.1101/153056

**Authors:** Zheng Liu, Maro J. Kariya, Christopher D. Chute, Amy K. Pribadi, Sarah G. Leinwand, Ada Tong, Kevin P. Curran, Neelanjan Bose, Frank C. Schroeder, Jagan Srinivasan, Sreekanth H. Chalasani

## Abstract

Animals respond to predators by altering their behavior and physiological states, but the underlying signaling mechanisms are poorly understood. Using the interactions between *Caenorhabditis elegans* and its predator, *Pristionchus pacificus*, we show that neuronal perception by *C. elegans* of a predator-specific molecular signature induces instantaneous escape behavior and a prolonged reduction in oviposition. Chemical analysis revealed this predator-specific signature to consist of a class of sulfolipids, produced by a biochemical pathway required for developing predacious behavior and specifically induced by starvation. These sulfolipids are detected by four pairs of *C. elegans* amphid sensory neurons that act redundantly and recruit cyclic nucleotide-gated (CNG) or transient receptor potential (TRP) channels to drive both escape and reduced oviposition. Specific abolishment of predator-evoked *C. elegans* responses by the anti-anxiety drug sertraline as well as functional homology of the delineated signaling pathways suggests a conserved or convergent strategy for managing predator threats.

## Introduction

Animal survival depends on the ability to sense predators and generate specific behavioral responses, such as flight or freezing. Examples from multiple vertebrate and invertebrate species indicate that prey use multiple sensory modalities (including vision, audition, and most frequently olfaction) to detect predators^1^^-^^3^. In particular, the chemosensory neurons in the rodent vomeronasal organ (VNO), Grueneberg ganglion, and main olfactory epithelium have been shown to have the ability to detect chemical signals from cat urine and generate stereotyped defensive behaviours^4,5^. The nature of these sensory circuits and signaling pathways that drive these invariant defensive behaviors, however, have remained elusive.

We approached this question by analyzing the behavioral responses of the nematode *Caenorhabditis elegans*^6^ to a predatory nematode *Pristionchus pacificus*^7^. These two nematodes likely shared a common ancestor around 350 million years ago^8^. Recent studies have shown that *P. pacificus* is a facultative predator. *P. pacificus* can bite and kill *C. elegans*^9^, a process facilitated by the extensive re-wiring of the *P. pacificus* nervous system under crowded and/or starvation conditions^10^. The nematode *C. elegans,* with its fully mapped neural network comprising of just 302 neurons connected by identified synapses^11^ and powerful genetic tools, is ideally suited for a molecular and circuit-level analysis of complex behaviors^12-15^. Combining chemical and genetic methods, we dissected the signaling circuits underlying *C. elegans’* responses to P. pacificus. We found that a novel class of sulfated small molecules excreted by *P. pacificus* trigger defensive responses in *C. elegans*. These *P. pacificus-*derived chemical signals are detected by *C. elegans* via multiple sensory neurons and processed via conserved neurotransmitter signaling pathways. Our results suggest that signaling pathways that process predator threats are likely conserved between *C. elegans* and more complex animals.

## Results

### *C. elegans* generates rapid avoidance and reduced egg laying in response to a predator

*C. elegans* was originally isolated from compost heaps in the developmentally arrested dauer stage^16,17^. However, recent studies have isolated proliferating and feeding populations of *C. elegans* from rotting flowers and fruits^17^^-^^19^, where they are often found to cohabit with other nematodes including the Diplogastrid *Pristionchus*^16^. Previous reports have shown that the terrestrial nematode, *P. pacificus* can kill and consume the smaller nematode *C. elegans*^9^. Further, when these two nematodes are placed together on an agar plate, *C. elegans* rapidly avoids *P. pacificus* (data not shown). We hypothesized that the prey, *C. elegans*, detects the predator, *P. pacificus*, through chemical cues and thus tested *C. elegans* responses to *P. pacificus* excretions. We found that *C. elegans* showed immediate avoidance behavior upon perceiving excretions of starved, but not well-fed predators (Fig. 1a and Supplementary Fig. S1a). Based on this result, we collected excretions from *P. pacificus* after 21 hours of starvation (“predator cue”) and found that it consistently repelled genetically diverse *C. elegans* isolates (Supplementary Fig. S1b). Chemotaxis assays indicated that these *C. elegans* avoidance behaviors were not in response to volatile components of predator cue (Supplementary Figs. S1c, S1d).

**Figure 1.**
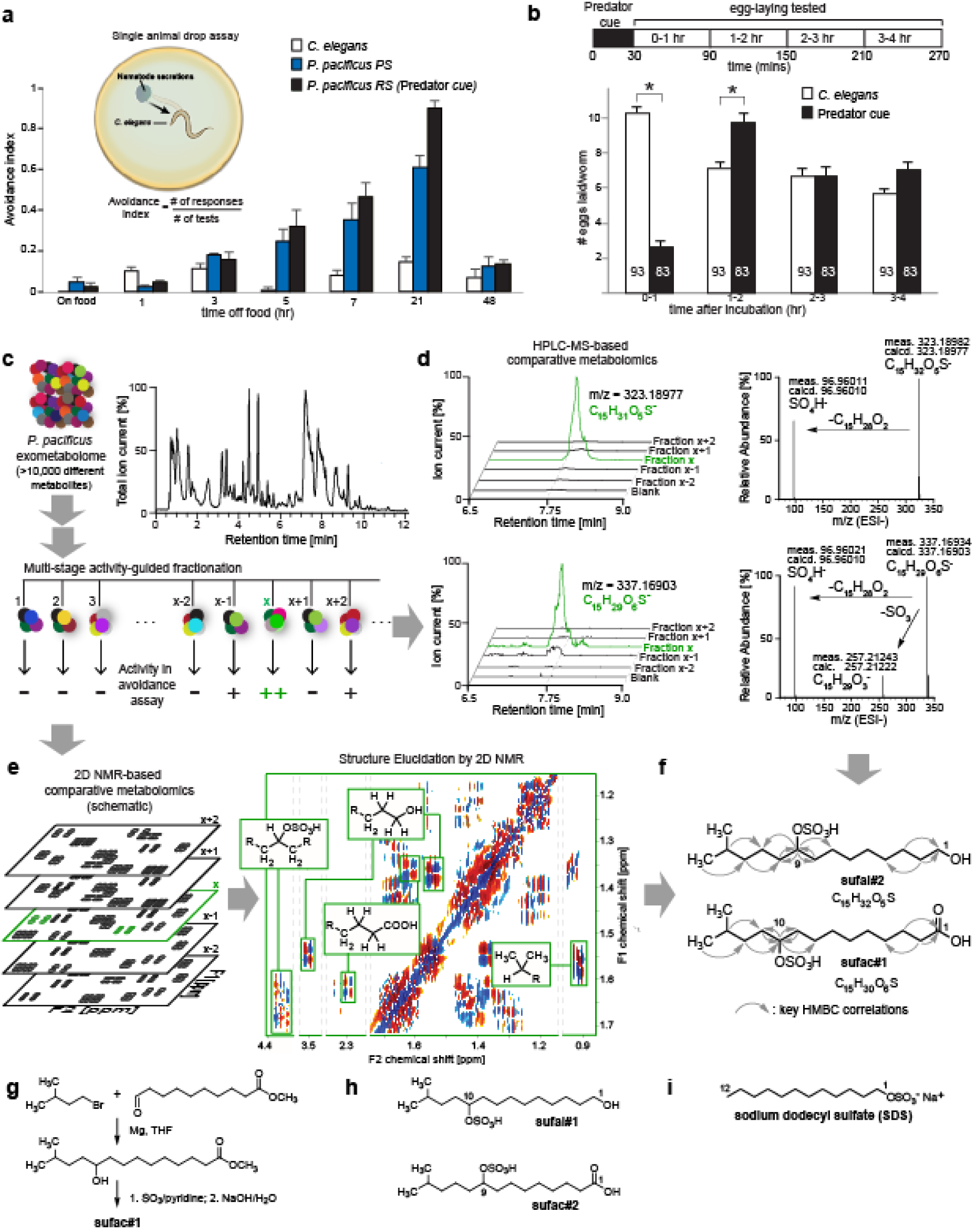
Predator-released sulfolipids drive *C. elegans* behaviors. **(a)** *C. elegans* avoid excretions from starving *P. pacificus* PS (PS312, domesticated) and RS (RS5725B a wild isolate, predator cue) strains. Inset, shows a schematic of the avoidance assay. **(b) Top** shows schematic of the modified egg laying assay and **bottom**, *C. elegans* lay fewer eggs after a 30-minute exposure to concentrated predator cue, but recovers after 2 hours. **(c)** UHPLC-HRMS analysis reveals a complex mixture of more than 10,000 metabolites, which was subjected to multistage activity-guided fractionation using reverse-phase chromatography. After four fractionation steps, most of the activity (++) was found in fraction x. Averages and s.e.m. are shown. n > 90 for each condition. **(d)** UHPLC-HRMS ion chromatograms (m/z value ± 5 ppm) of active fraction x and adjacent fractions for two sulfate containing metabolites that were strongly enriched in the active fraction (left). MS-MS analysis (right) confirms presence of sulfate moieties in both compounds. **(e)** Schematic representation of 2D NMR-based comparative metabolomics (left) of consecutive fractions (x-2 to x+2) used to identify signals specific to fraction x. Cropped 2D NMR (dqfCOSY) spectrum (right) of active fraction highlighting signals that represent specific features of the identified metabolites (grey lines define edges of shown subsections). **(f)** Chemical structures of metabolites identified via comparative metabolomics from active fraction x, sufac#1 and sulfal#2. Grey arrows indicate important correlations observed in heteronuclear 2D NMR (HMBC) spectra. **(g)** Synthesis of sufac#1; THF: tetrahydrofuran. **(h)** Homologous metabolites sufac#2 and sufal#1 were also detected by UHPLC-HRMS. **(i)** Chemical structure of sodium dodecyl sulfate (SDS). Averages and s.e.m. are shown and number of animals tested are indicated on each bar or condition. *p < 0.05 obtained by comparison with controls using Fisher’s exact t-test with Bonferroni correction.

We further found that *C. elegans* exposed to predator cue did not lay eggs for many minutes following exposure, even when placed on food (bacterial lawn), suggesting that predator cue-induced stress affects egg laying behavior. Consistent with this idea, previous studies have shown that *C. elegans* retain eggs in the gonad when exposed to environmental stressors^20^. To test our hypothesis, we designed a behavioral assay wherein the prey was exposed to predator cue for 30 minutes, and egg laying was monitored for many hours following cue removal. Animals exposed to predator cue laid significantly fewer eggs than controls during the initial 60 minutes following cue removal. During the next hour (i.e., the 60–120-minute post-cue time period), these animals laid more eggs than controls, suggesting that predator cue transiently modified egg laying behavior, but not egg production (Fig. 1b). Collectively, these results indicate that starving *P. pacificus* release a potent, non-volatile factor (predator cue) that elicits multiple prey responses, namely urgent escape behavior followed by up to one hour of reduced egg laying.

### Predator releases a novel family of sulfolipids to drive *C. elegans* responses

We aimed to identify chemical structure(s) of the small molecule(s) excreted by *P. pacificus* that caused *C. elegans* avoidance behavior. Because the *P. pacificus* exo-metabolome is highly complex, consisting of more than 20,000 distinct compounds detectable by UPLC-HRMS (Fig. 1c), we used a multi-stage activity-guided fractionation scheme (see Supplementary Methods). After three rounds of fractionation, comparative analysis of 2D NMR spectra^21,22^ and high-resolution tandem mass spectrometry data of active and adjacent inactive fractions (Figs. 1d, 1e, Supplementary Tables S1-S3), revealed several novel (ω-1)-branched-chain sulfolipids (sufac#1, sufac#2, sufal#1, and sufal#2) as major components of active, but not inactive fractions (Fig. 1f). We then synthesized these compounds and tested their activity in the avoidance assay. The terminal alcohols sufal#1 and sufal#2 accounted for most of the isolated activity (Fig. 1g, Supplementary Fig. S3a, Supplementary methods). Following identification of these compounds, we re-analyzed the *P. pacificus* exo-metabolome by UPLC-MS/MS, which revealed a large number of structurally related sulfolipids (Fig. 1h). None of the *P. pacificus* sulfolipids could be detected in the metabolomes of *E. coli* (used as food for nematodes) (Supplementary Fig. S2), *C. elegans*, or several other nematode species (Supplementary Table S4), which were extracted and analyzed under identical conditions. Notably, the identified sulfolipids are structurally similar to sodium dodecyl sulfate (SDS, Fig. 1i), which is a potent *C. elegans* avoidance cue ^23^. Additional assays showed that the sulfolipids identified from *P. pacificus* excretions also attenuated *C. elegans* egg laying (Supplementary Fig. S3b), demonstrating that these predator-specific small molecules instruct both rapid and longer-lasting prey responses.

### Multiple *C. elegans* sensory neurons act redundantly to generate predator avoidance

To define the prey neural circuit that detects predator cue, we tested the role of all 12 pairs of amphid sensory neurons, which project dendrites to the nose of the animal to sense environmental changes (Fig. 2a)^11,24^. Previous studies have shown that sensory neurons in the amphid ganglia located in the head of the worm detect repellents and generate reversals in an attempt to avoid the noxious cues^24,25^. We found that animals lacking pairs of ASJ, ASH, ASI, or ADL neurons (but not other amphid neurons) were defective in their responses to predator cue (Figs. 2b, 2c), indicating that *C. elegans* uses multiple sensory neurons to detect predators. Responses to sulfolipids purified from predator cue and SDS were similarly reduced in animals lacking these neurons (Supplementary Figs. S3f-g).

**Figure 2.**
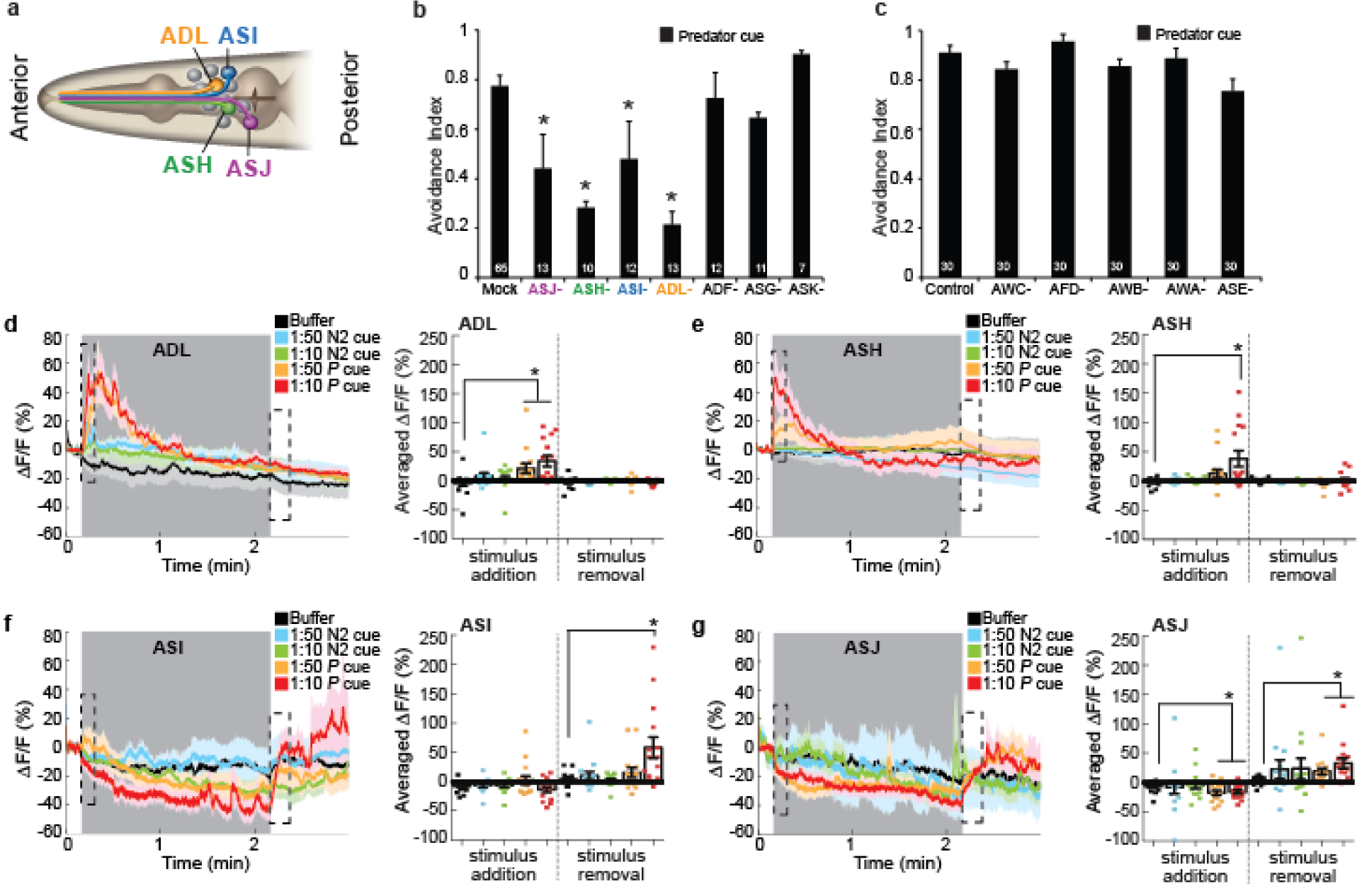
Multiple sensory neurons are required for avoidance of predator cue. **(a)** Schematic showing amphid sensory neurons with key neurons highlighted. **(b)** Cell and **(c)** genetic ablations showing that ASJ, ASH, ASI, and ADL sensory neurons, but not other amphid neurons, are required for avoidance of predator cue. Averages and s.e.m. and numbers of animals tested are shown on each bar. **(d-g)** Average calcium responses of transgenic animals (n > 13 for each condition) expressing the GCaMP family of indicators in **(d)** ADL, **(e)** ASH, **(f)** ASI, or **(g)** ASJ sensory neurons to predator cue (*P* cue) or *C. elegans* secretions (N2 cue). Each experiment was a 180 second recording where control (M9 buffer), *C. elegans* secretions (N2 cue), or predator cue (*P* cue) in different dilutions was added at 10 seconds and removed at 130 seconds (stimulus is indicated by a shaded grey box). Bar graphs, average percentage change during the 10 seconds after stimulus addition (dashed box), or 15 seconds after stimulus removal (dashed box) are shown. Error bars and shaded regions around the curves represent s.e.m. *p < 0.05 obtained by comparison with controls using Fisher’s exact t-test with Bonferroni correction.

To confirm the role for ADL, ASH, ASI, and ASJ neurons, we monitored their responses to predator cue using calcium imaging^14^. Calcium responses are strongly correlated with neuronal activity in *C. elegans* neurons ^26,27^. We found that adding predator cue to the nose of the prey activated ADL and ASH (Figs. 2d, 2e, Supplementary Fig. S4 for all traces), whereas predator cue removal activated ASI and ASJ neurons (Figs. 2f, 2g, Supplementary Fig. S4 for all traces). Also, whereas ADL and ASJ responded to both tested dilutions of predator cue (Figs. 2d, 2g), ASH and ASI only detected the more concentrated cue (i.e., 1:10 dilution, but not 1:50) (Figs. 2e, 2f), suggesting different response thresholds for these four neuronal pairs. Collectively, these results show that predator cue activates ADL and ASH neurons, whereas its removal increases ASI and ASJ activity.

### Sensory neurons use CNG and TRP channels to drive predator avoidance

To gain insight into the signal transduction machinery underlying these responses, we examined the behavior of mutants lacking specific signaling components. We found that mutants lacking the alpha subunit (*tax-4*), but not the beta subunit (*tax-2*), of the cyclic nucleotide gated (CNG) ion channel exhibited defective responses to predator cue (Fig. 3a). Moreover, expressing the full-length *tax-4* cDNA via a *tax-4* promoter, an ASI-specific promoter, or an ASJ-specific promoter (but not via an ASH-selective promoter) restored normal behavior to the null mutants (Fig. 3a). The ability of these transgenes to rescue avoidance behavior was largely dose dependent, as it varied depending on the amount of *tax-4* transgene expressed in ASI and ASJ neurons (Supplementary Fig. S5a). These data indicate that increased CNG signaling from ASI neurons could compensate for the lack of signaling from ASJ, and vice-versa. Similarly, mutants lacking the transient receptor potential (TRP) channel OCR-2, but not OSM-9, were defective in their responses to predator cue. Further, we observed that OCR-2 functions in ADL and ASH neurons, but not in ASI or ASJ neurons (Fig. 3b), and that responses to ASH- and ADL-specific *ocr-2* transgenes were also dose dependent (Supplementary Fig. S5b), indicating that signaling from ASH could compensate for the lack of ASJ signaling, and vice-versa. Testing samples of purified sulfolipids confirmed that *tax-4* (but not *tax-2*) and *ocr-2* (but not *osm-9*) mutants were defective in avoidance to these molecules (Supplementary Fig. S5c). Also, since TAX-4 but not TAX-2 can form a homomeric CNG channel^28^, we hypothesize that the OCR-2 TRP channel may either form a homomeric channel that interacts with other non-OSM-9 TRP channel subunits to generate a functional channel and drive avoidance behavior, results consistent with previous studies^29,30^.

**Figure 3.**
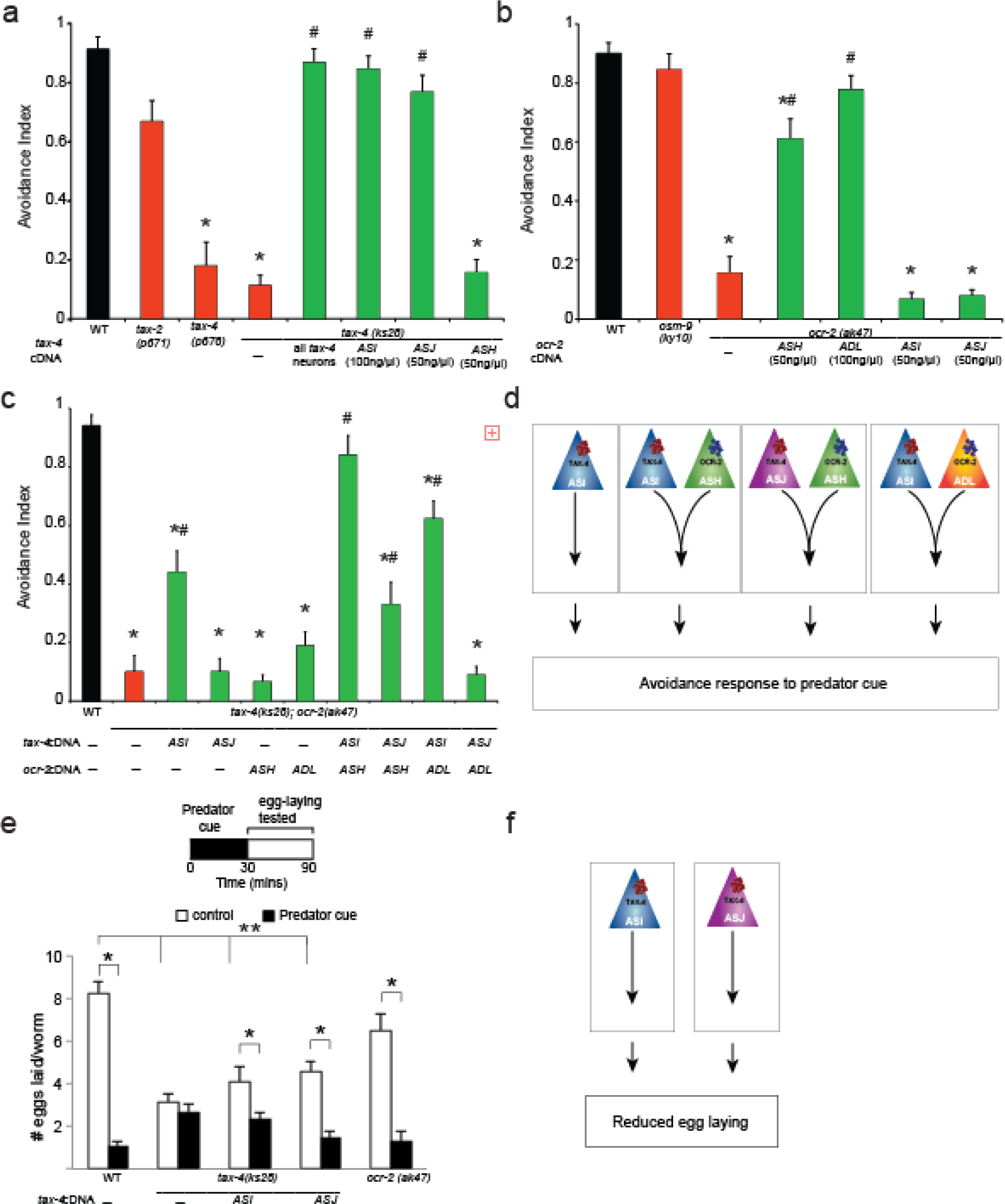
A redundant *C. elegans* signaling network enables responses to predator cue. **(a)** Mutants lacking the alpha subunit of the CNG channel (*tax-4)*, but not the beta subunit (*tax-2),* are defective in their response to predator cue. Restoring wild-type *tax-4* cDNA using an ASI- or ASJ-specific promoter is sufficient to restore normal behavior to *tax-4* mutants. **(b)** Mutants lacking the TRPV channel subunit *ocr-2*, but not *osm-9,* are defective in their response to predator cue. OCR-2 is specifically required in ASH sensory neurons. **(c)** A *tax-4; ocr-2* double mutant is also defective in avoiding predator cure and wild-type behaviour is restored when TAX-4 is restored to ASI at the same time OCR-2 is restored to ASH neurons. Restoring TAX-4 to ASJ and OCR-2 to ASH or ADL or restoring TAX-4 to ASI and OCR-2 to ADL is able to partially restore wild-type behaviour to the double mutants. Moreover, expressing *tax-4* in ASI but not ASJ is sufficient to rescue of the double mutant phenotype. **(d)** *C. elegans* egg-laying reduction requires functional TAX-4 signaling in ASI and ASJ neurons. OCR-2 is not required for this behaviour. Schematic showing **(e)** four redundant pathways (ASI acting independently, ASI and ASH, or ASI and ADL, or ASJ and ASH acting together) driving avoidance to the *P. pacificus* predator cue. **(f)** Schematic showing that TAX-4 acts in ASI and/or ASJ neurons to reduce egg laying upon longer-term exposure to predator cue. Averages and s.e.m. are shown. (a-c) n = 90 animals and in **(e)** n > 35 tested for each condition, *p < 0.05 compared with controls, ^#^p < 0.05 compared to mutants, **p < 0.05 comparing eggs laid by mutants and wild type animals obtained using Fisher’s exact t-test with Bonferroni correction.

To investigate possible interactions between CNG and TRP channel signaling, we analyzed *tax-4; ocr-2* double mutants. Restoring TAX-4 function to ASI neurons and OCR-2 to ASH neurons (in combination) conferred normal predator cue avoidance to the double mutants (Fig. 3C). Moreover, partial avoidance was seen for other rescue combinations (TAX-4 in ASJ and OCR-2 in ASH; TAX-4 in ASI and OCR-2 in ADL, and TAX-4 in ASI alone) (Fig. 3C). Together, these data indicate that there are at least four neuronal signaling pathways that can drive robust avoidance to *Pristionchus* cue: 1) ASI sensory neurons using TAX-4 channels, 2) ASI and ASH neurons using TAX-4 and OCR-2 channels, respectively, 3) ASJ and ASH using TAX-4 and OCR-2 channels, respectively, and 4) ASI and ADL using TAX-4 and OCR-2 channels, respectively (Figs. 3d). Similarly, we found that *tax-4* mutants, but not *ocr-2* mutants, did not curtail their egg laying behavior (a longer-lasting effect) in response to predator cue, and that restoring TAX-4 function to ASI or ASJ significantly improved this *tax-4* defect (Figs. 3e, 3f).

### Sertraline acts on GABA signaling to block predator-evoked *C. elegans’* responses

Next, we asked whether *C. elegans* avoidance behavior could be suppressed by small molecules that alleviate mammalian anxiety (Supplementary Table S5)^31,32^. We screened a small library of anti-anxiety drugs, and found that pre-treating prey with a selective serotonin reuptake inhibitor, sertraline (brand name “Zoloft”) attenuated avoidance to predator cue and purified sulfolipids, but not to other repellents (Fig. 4a, Supplementary Table S5). Suppression of avoidance behavior by sertraline was dose-dependent (Supplementary Fig. S6a) and lasted for at least 30 minutes after the drug was removed (Supplementary Fig. S6b). To test whether sertraline modifies signaling from specific sensory neurons, we analyzed mutants expressing different rescuing transgenes. Sertraline had no detectable effect on the behavior of *tax-4* or *ocr-2* mutants, but it attenuated avoidance to predator cue of: 1) *tax-4* mutants expressing *tax-4* in ASI or ASJ, and 2) *ocr-2* mutants expressing *ocr-2* in ADL or ASH (Fig. 4b). Thus, sertraline affected signaling from all four sensory neurons. Next, we found that sertraline required GABA, but not serotonin signaling. Animals lacking glutamic acid decarboxylase (*unc-25*, enzyme required for GABA synthesis^33^), but no other neurotransmitter biosynthetic enzymes were defective in sertraline attenuation (Fig. 4c, Supplementary Fig. S6c). Additionally, adding GABA exogenously to the agar plate was sufficient to restore normal behavior to *unc-25* mutants confirming that GABA signaling is required to modify predator avoidance (Fig. 4c). Further, we found that restoring UNC-25 function to all 26 GABAergic neurons^34^ or under a RIS interneuron-selective promoter was sufficient to restore sertraline attenuation of predator avoidance (Fig. 4d). RIS interneuron has been previously shown to play a role in inducing a sleep-like state in *C. elegans*^35,36^ and these results suggest an additional for this neuron in modifying predator behavior. Moreover, we found that mutants in a solute carrier 6 plasma membrane re-uptake transporters [*snf-10* ^37^], but no other GABA transporters, were partially defective in their response to predator cue after sertraline treatment (Fig. 4e). Consistently, rodent and human homologs of this protein have been shown to bind multiple SSRIs using a non-competitive mechanism^38^. Finally, we found that sertraline treatment also reduced the longer-lasting egg laying response (Fig. 4f) showing that the drug blocks *C. elegans’* responses on multiple timescales. Taken together, these results indicate that the anti-anxiety drug sertraline specifically abolished predator-induced *C. elegans* responses by acting on GABA signaling in RIS interneuron.

**Figure 4.**
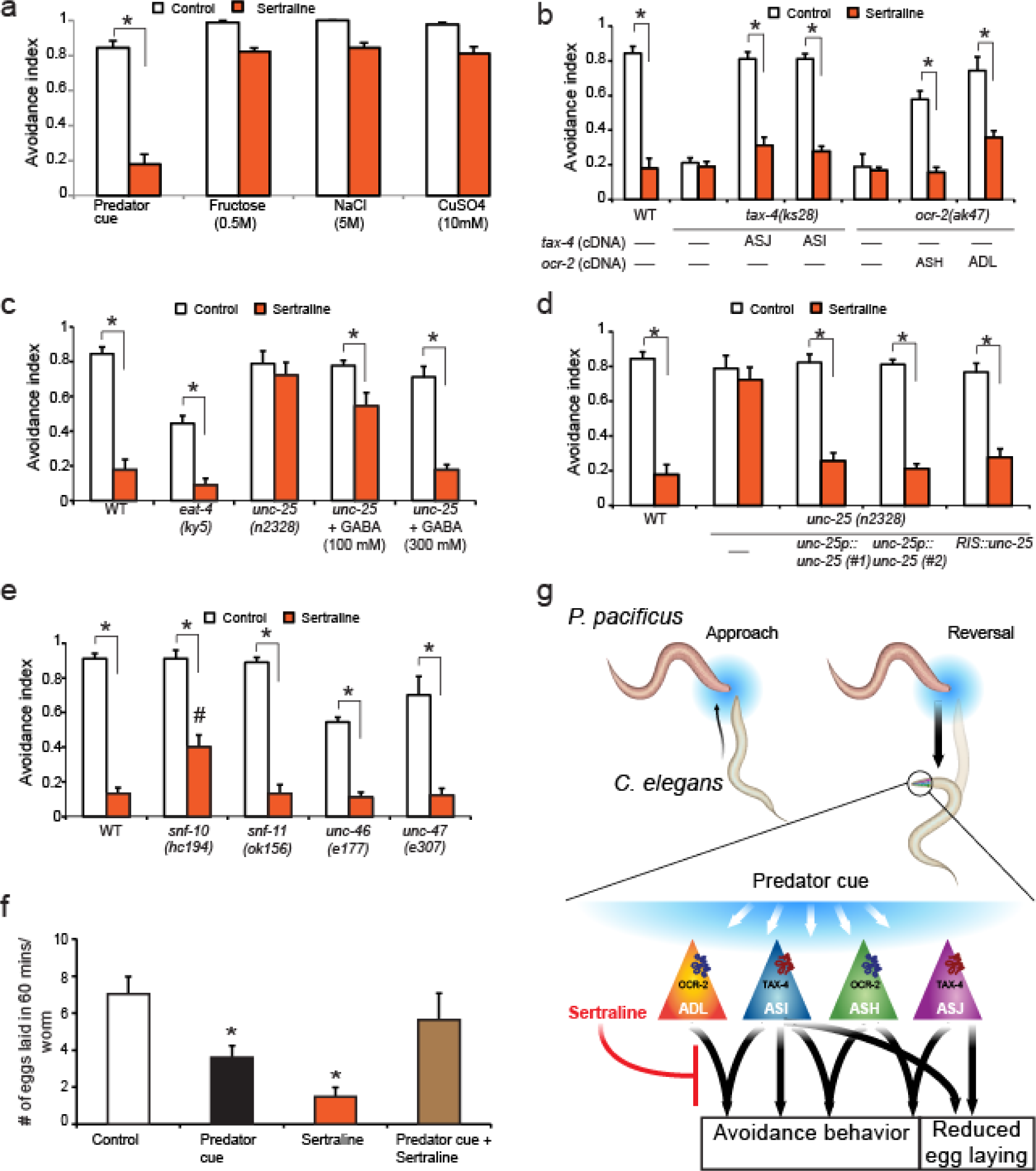
Sertraline attenuates *C. elegans* responses to the *P. pacificus* predator. **(a)** Sertraline specifically attenuates *C. elegans* responses to predator cue, but not to fructose, salt, or copper. **(b)** The effect of sertraline is lost in *tax-4* and *ocr-2* mutants, and modulation is restored when TAX-4 is restored to either ASI or ASJ, or when OCR-2 is restored to ASH or ADL. **(c)** Sertraline requires GABA, but not glutamate signaling. Sertraline modulates avoidance responses of *eat-4* (glutamate receptor), but not *unc-25* (glutamic acid decarboxylase, required for GABA synthesis) mutants. Adding GABA exogenously to plates is able to restore sertraline modulation to *unc-25* mutants. **(d)** Defective *unc-25* response is rescued by expressing wild-type *unc-25* cDNA under the *unc-25* promoter or a RIS-selective promoter. **(e)** Sertraline partially modulates mutants in a plasma membrane transporter (*snf-10*) and completely attenuates mutants in other GABA transporters, *unc-47*, *snf-11*, and *unc-46* (a transmembrane protein that recruits UNC-47). **(f)** Animals were treated with predator cue or 1 mM sertraline or predator cue and sertraline for 30 minutes and egg-laying was monitored for 60 minutes after removal of these compounds. **(g)** *C. elegans* detects secretions from a starving *P. pacificus* (predator cue) using sensory circuits consisting of ASI, ASJ, ASH, and ADL neurons that use CNG and TRP channels and act in a redundant manner to generate rapid avoidance. In contrast, CNG channels act in ASI and ASJ neurons to reduce egg laying for many minutes. Sertraline attenuates both predator-induced avoidance behaviour and egg laying behaviour downstream of these sensory neurons. Averages of either n > 90 **(a-e)** or n > 35 **(f)** and s.e.m. are shown. *p < 0.05 compared to controls obtained using Fisher’s exact t-test with Bonferroni correction

## Discussion

We show that *P. pacificus* releases a mixture of sulfolipids that *C. elegans* perceives as a predator-specific molecular signature, or kairomone^39^. Perception of these sulfolipids via multiple sensory neurons initiates a fear-like avoidance response and reduces egg laying behavior (Fig. 4f). Among the nematode species whose metabolomes we have analysed, *P. pacificus* is the only one that excretes copious amounts of sulfolipids. Sulfated fatty acids and related lipids have previously been described primarily from marine sources, including tunicates^40^, sponges^41^, crustaceans^42^, and algae^43^. In addition, a family of sulfated fatty acids, the caeliferins, has been identified from grasshopper oral secretions^44,45^. In a striking parallel to the role of sulfated lipids in the nematode predator-prey system studied here, these herbivore-associated sulfolipids have been shown to elicit specific defense responses in plants^45^. Furthermore, the sulfolipids we identified from *P. pacificus* resemble sodium dodecyl sulfate (SDS), a known nematode repellent^23^. We found that, similar to avoidance triggered by predator cue, ASJ, ASH and ASI neurons are necessary for avoidance to SDS (Supplementary Fig. S3g). Given the similarity of the neuronal circuitry required for the avoidance responses, it appears that *C. elegans* avoid SDS because of its structural similarity to the *Pristionchus*-released sulfates, which are interpreted as a molecular signature of this predator.

The sulfolipids we identified from *P. pacificus*, sufac#1 and sufal#2, and several related compounds, appear to be derived from the monomethyl branched-chain fatty acid (mmBCFA), C15ISO, which is also produced by *C. elegans* and has been shown to be essential for *C. elegans* growth and development^46^. The biosynthesis of C15ISO in *C. elegans* requires the fatty acid elongase ELO-5, and several homologous elongases in *P. pacificus* exist that may be involved in the biosynthesis of the fatty acid precursors of sufac#1 and sufal#2. Additionally, the biosynthesis of sufac#1 and sufal#2 requires oxygenation at the (ω-5) or (ω-6) position in the fatty acid chain, respectively, followed by sulfation by sulfotransferase(s), a family of genes that has undergone major expansion in *P. pacificus*^47^. Notably, at least one sulfotransferase, EUD-1, functions as a central switch determining whether *P. pacificus* larvae will develop into a primarily bacterivorous, narrow-mouthed adult, or into a predacious, wide-mouthed adult that can feed on other nematodes^48^. It is intriguing that *C. elegans* has evolved the ability to detect a *Pristionchus*-specific trait (the extensive sulfation of small molecules) that is directly connected to the endocrine signaling pathway that controls development of the morphological features required for predation.

Detection of predator cue relies on a sensory neural circuit consisting of at least four different amphid neurons (ASI, ASH, ASJ and ADL, Fig. 4g). These neurons have well-described roles in detecting chemicals from the environment: the ASI and ASJ sensory neurons play a major role in the detection of ascaroside pheromones, while ASH neurons are nociceptive and drive avoidance to glycerol and copper, and ASH and ADL act together to promote social feeding ^23,49-53^. Therefore, whereas ASH and ADL have been shown to drive avoidance behavior^23,52,54^, our finding that ASI and ASJ are involved in generating avoidance are novel. Participation of these additional neurons facilitates redundancy, such that signaling from ASI, ASI and ASH, ASI and ADL, ASJ and ASH, and ASJ and ADL is sufficient to drive avoidance to predator cue. Similarly, signaling from either ASI or ASJ neurons alters egg-laying behavior. Such redundant circuit(s) are likely to decrease the failure rate for signaling, thereby increasing the robustness of the behavioral output. Similar redundant circuits have been described for sensory neurons detecting temperature^55^ or odors^56^, and in neural circuits driving feeding in the crab^57^.

The neuronal signaling machinery in the ASI, ASJ, ASH and ADL sensory neurons relies on CNG and TRP channels to mediate responses to predator cue. CNG ion channels typically consist of alpha and beta subunits and have been shown to play a central role in regulating chemosensory behaviors across multiple species ^58,59^. Because the alpha subunit homolog TAX-4 is required for detecting predator cue, whereas the beta subunit TAX-2 is not, we suggest that homomeric TAX-4 channels act in the ASI and ASJ sensory neurons. *In vitro* experiments have shown that *C. elegans* TAX-4 subunits can form a functional homomeric channel when expressed in HEK293 cells^60^. Similarly, alpha subunits of the CNG channels have also been shown to function as homomeric channels both *in vitro* and *in vivo*^61,62^. Our studies also indicate a role for a subunit of the TRP channel OCR-2, but not its heteromeric partner OSM-9^63^. We suggest that OCR-2 can either form a homomeric channel or interact with other non-OSM-9 TRP channel subunits to generate a functional channel and drive avoidance behavior. These results are consistent with previous studies where OCR-2 has been shown to act independently of OSM-9 in regulating *C. elegans* larval starvation^29^ and egg-laying behaviors^30^. Our results for the role of TRP channels are reminiscent of rodent studies where TRP channels have been found to play a crucial role in initiating responses to predator odors from cats^4^, suggesting broad conservation of the molecular machinery that detects predators. Taken together, we hypothesize that signaling from homomeric CNG and TRP channels acting in distinct, but redundant sensory circuits enables reliable detection of predators by the prey.

We further show that the anti-anxiety drug, sertraline, acts downstream of CNG and TRP channels and requires GABA signaling in RIS interneurons to suppress predator-evoked responses. Sertraline has been shown to particularly effective in alleviating human anxiety disorders^64^ and, classified as an SSRI, is thought to act in part by elevating serotonin levels at synapses^65,66^. Our studies show that sertraline requires GABA, but not serotonin signaling, to exert its effects on *C. elegans* avoidance behavior. Other SSRIs have also been shown to require GABA signaling in mammalian^67,68^ and *C. elegans* nervous systems^69^, in addition to effects on other neurotransmitter pathways including dopamine, glutamate, histamine, and acetylcholine^70^. We further show that sertraline action in *C. elegans* requires functional glutamic acid decarboxylase, a GABA biosynthesis enzyme^33^ specifically in RIS interneurons, defining the site of action of the drug. RIS interneurons have been implicated in modulating a sleep-like state in *C. elegans*^35,36^. We also found that sertraline effects require a plasma membrane re-uptake transporter that belongs to the solute carrier 6 family (*snf-10*). This protein has been previously implicated in activating *C. elegans* sperm in response to male protease activation signals^37^. While our results showing a role for *snf-10* in animal behavior are novel, the closely related *snf-11* has been shown to act to clear GABA from synaptic clefts^71^. We suggest that sertraline might act on the SNF-10 protein at RIS synapses to modify *C. elegans* behavior. This idea is consistent with results from *in vitro* studies where rodent and human homologs of this protein have been shown to bind multiple SSRIs using a non-competitive mechanism^38^. Collectively, our analysis of the mechanisms by which sertraline attenuates *C. elegans* avoidance responses to an external stressor (predator) suggest broad conservation of the involved signaling pathways from worms to humans.

In summary, our results uncover the chemosensory and neuronal basis of a predator-prey relationship between *P. pacificus* and *C. elegans*, in which predator detection is based on a characteristic molecular signature of novel sulfate-containing molecules. The prey uses multiple sensory neurons acting in parallel and conserved CNG and TRP channel signaling to detect these sulfates and drive rapid avoidance and longer lasting reduced egg laying. Additionally, we show that sertraline acts on GABA signaling in RIS interneurons likely targeting a plasma membrane re-uptake transporter to attenuate *C. elegans* avoidance behavior. Based on these results, we hypothesize that *C. elegans* evolved mechanisms to detect *Pristionchus*-released sulfolipids as a kairomone, and that the identified neuronal signaling circuitry is representative of conserved or convergent strategies for processing predator threats.

## Methods Summary

Single animal avoidance assay and calcium imaging were performed as described^14,23^. Avoidance indices of all strains tested and controls are shown in Supplementary Table S6, while all the neuronal responses are shown in Supplementary Fig. S4.

Egg laying assays were modified from assays previously described^20^, where synchronized adults were exposed to test compound for 30 minutes and the number of eggs laid monitored for the times indicated.

Additional methods and sulfolipid purification and synthesis are shown in supplementary information online.

## Supplementary Information

Supplemental information includes 6 figures and 7 tables, methods, and NMR spectra.

## Acknowledgements

We thank R. Hong, R. Sommer, C. Bargmann, S. Mitani, the National BioResource Project and Caenorhabditis Genetics Center (CGC) for strains; C. Bargmann and W. Liedke for constructs; and P. Sengupta, A. Chisholm, R. Hong, C. Bargmann, Y. Jin, D. O’Keefe, K. Quach, U. Magaram, H. Lau, L. Hale, and other members of the Chalasani Laboratory for helpful discussions and comments on the manuscript. This work was supported by grants from the W. M. Keck Foundation, NIH R01MH098001, R01MH113905 (S.H.C.), NIH R01AT008764 (F.C.S.), a Salk Alumni Fellowship (K.P.C) and Startup Funds from Worcester Polytechnic Institute (J.S.).

## Author Contributions

Z.L., M.J.K., C.D.C., A.K.P., S.G.L., A.T., K.P.C., and N.B. performed experiments, developed experimental methods and reagents. F.C.S., J.S., and S.H.C. designed and interpreted the experiments and wrote the paper.

